# QMAP: A Benchmark for Standardized Evaluation of Antimicrobial Peptide MIC and Hemolytic Activity Regression

**DOI:** 10.64898/2026.02.03.703041

**Authors:** Anthony Lavertu, Jacques Corbeil, Pascal Germain

## Abstract

Antimicrobial peptides (AMPs) are promising alternatives to conventional antibiotics, but progress in computational AMP discovery has been difficult to quantify due to inconsistent datasets and evaluation protocols. We introduce QMAP, a domain-specific benchmark for predicting AMP antimicrobial potency (MIC) and hemolytic toxicity (HC50) with homology-aware, predefined test sets. QMAP enforces strict sequence homology constraints between training and test data, ensuring that model performance reflects true generalization rather than overfitting. Applying QMAP, we reassess existing MIC models and establish baselines for MIC and HC50 regression. Results suggest limited progress over six years, poor performance for high-potency MIC regression, and low predictability for hemolytic activity, emphasizing the need for standardized evaluation and improved modeling approaches for highly potent peptides. We release a Python package facilitating practical adoption, and with a Rust-accelerated engine enabling efficient data manipulation, installable with `pip install qmap-benchmark`.

## Introduction

The discovery of antibiotic agents in the twentieth century was one of the most important discoveries of modern medicine. Unfortunately, the abusive use of antibiotics pressured bacteria to develop resistance. Many bacteria have developed resistance to multiple classes of antibiotics, making them a threat to human health. If no solution is found, the damages caused by AMR could reach 100 billion dollars and 10 million deaths annually in 2050^1^.

In this context, antimicrobial peptides (AMPs) have attracted increasing attention as potential alternatives or complements to small-molecule antibiotics. AMPs are naturally synthesized by most living organisms as part of their innate immune defense mechanisms. A defining feature of AMPs is that they have co-evolved with bacteria over millions of years in a continuous evolutionary arms race, in which bacterial resistance mechanisms are countered by diversification and optimization of host peptides^2^. This long co-evolutionary history has resulted in a rich and diverse repertoire of AMPs and is associated with a slower emergence of resistance compared to conventional antibiotics^3^. Moreover, they have a set of properties suitable for clinical development, such as broad spectrum activity and high selectivity, which reduce toxicity^4^. These characteristics make AMPs a promising direction for future antibiotic research^5^, particularly as synergistic agents used in combination with classical antibiotics to enhance efficacy and mitigate resistance^6,7^.

The search for AMPs that are both highly potent and minimally toxic has motivated the adoption of machine learning and deep learning approaches to accelerate discovery and reduce experimental costs through in silico screening and rational peptide design. Within this framework, two quantitative measurements are of central importance: the Minimum Inhibitory Concentration (MIC) and the Hemolytic Concentration 50 (HC50). MIC measures the lowest concentration of a peptide required to inhibit bacterial growth and serves as a direct indicator of antimicrobial potency, with lower values corresponding to greater activity^8,9^. In contrast, HC50 quantifies the concentration at which a compound lyses 50% of erythrocytes and is therefore a measure of cytotoxicity, where lower values indicate higher toxicity. These two properties are typically in tension, as an increased antimicrobial potency can be accompanied by increased hemolytic activity. Consequently, AMP development aims to maximize potency while minimizing toxicity, which can be expressed as minimizing the ratio MIC*/*HC50.

The application of artificial intelligence to the modelling of antimicrobial peptides has expanded rapidly in recent years. In 2019, J. and Z. Witten introduced, to the best of our knowledge, the first deep neural network for predicting the MIC of peptides against E. coli, alongside the assembly of a dataset for MIC regression^10^. Building on this foundation, S. N. Dean et al. developed PepVAE, a variational autoencoder framework combining generative modelling with MIC regression to design potent antimicrobial peptides^11^. In 2022, S. Gull and F. Minhas proposed AMP0, a machine learning framework capable of predicting MIC values for arbitrary bacterial targets given access to the corresponding genome^12^. In 2023, J. Huang et al. introduced a multi-stage pipeline integrating classification, ranking, and regression models to explore the complete peptide space of lengths of 6–9 amino acids, identifying a peptide with therapeutic efficacy comparable to penicillin and negligible toxicity^13^. Shortly thereafter, J. Yan et al. presented a multi-branch convolutional neural network for MIC prediction against E. coli^14^. In 2024, Sharma et al. developed a recurrent neural network–based model targeting ESKAPEE pathogens^15^, and in the same year, C. Chung et al. proposed an ensemble combining recurrent, convolutional, multi-branch, and transformer architectures to predict MIC values for specific bacterial strains^16^. Most recently, in 2025, J. Cai leveraged transfer learning from transformer models to predict MIC values for E. coli and S. aureus^17^.

The development of models for quantitative prediction of hemolytic activity (HC50) is more recent. In 2025, A. S. Rathore et al. conducted a comparative evaluation of multiple algorithms and found that a protein language model achieved the highest correlation with experimentally measured hemolytic activity against mammalian erythrocytes^18^. Shortly thereafter, AMPLyze introduced a hybrid model combining embeddings from protein language models with sequence-level descriptors, optimized using a log-cosh loss function^19^. Later in 2025, X. Li et al. published HemPepPred, an ensemble approach integrating multiple tree-based models with ridge regression to predict HC50 values^20^.

Despite these methodological advances, quantitative comparison between models remains challenging. Reported test-set performance is highly sensitive to the particular choice of test data and to the random seed used during dataset splitting. Moreover, many studies rely on random or only weakly constrained splitting strategies, which can allow closely related peptide sequences to appear in both training and test sets. This practice likely contributes to the wide variation in reported results across studies, even when models are evaluated on datasets described as independent.

Similarity-aware splitting strategies that explicitly prevent information leakage in sequence space are already available. Notably, GraphPart^21^ formulates datasets splitting as a graph partitioning problem, ensuring that sequences exceeding a specified identity threshold cannot be assigned to different splits. DataSail^22^ represents a conceptually related approach that aims to generate splits that are both similarity-aware and balanced with respect to multiple dataset constraints. Despite their methodological rigour, these strategies have seen limited adoption in the AMP MIC and hemolytic regression literature. As a result, most studies continue to rely on random or weakly constrained splits, undermining the reliability of quantitative model comparisons. In parallel, the field has lacked a domain-specific benchmark tailored to AMP regression tasks, leading each study to define its own dataset, split strategy, and evaluation protocol. PepBenchmark^23^ represents an important step toward standardized evaluation in bioactive peptide research; however, its general-purpose design and reliance on user-defined splits limit its suitability for AMP-specific MIC regression, as different studies may evaluate models on different test sets. Furthermore, it does not provide a principled mechanism to incorporate external training data while preserving a controlled minimal homology distance between training and test sets, which is essential for fair comparison and realistic assessment of generalization.

To address these challenges, we introduce the Quantitative Mapping of Antibacterial Peptide benchmark (QMAP), a domain-specific benchmark for AMP MIC and hemolytic HC50 regression that provides predefined, homology-aware test splits, ensuring that all models are evaluated on exactly the same data. QMAP includes a protocol to verify and enforce a minimum homology distance between any user-defined training dataset and the benchmark test sets, enabling fair and reproducible comparison across studies. This verification is implemented using a graph-based similarity framework inspired by GraphPart^21^. Beyond this methodological rigor, a well-populated benchmark serves as a living showcase of the state of the art, with direct practical value for the field. Applied researchers can leverage top-performing models to screen large peptide libraries for candidate discovery or as reward signals in generative design pipelines, with promising candidates subsequently validated experimentally. More fundamentally oriented researchers can use these models as a starting point for knowledge extraction, interpretability studies, or distillation into mechanistically transparent models. While QMAP does not itself provide new peptide candidates or mechanistic insights, it provides a standardized foundation that facilitates such downstream work. Using this benchmark, we evaluate representative MIC regression models from the literature as well as a baseline approach for HC50 prediction. Our results provide evidence that progress in the field has been more limited than previously suggested, and the benchmark highlights key areas where future methodological advances are needed. To facilitate practical adoption, we release an open-source Python package alongside this work, providing tools for data manipulation, similarity-aware splitting, and access to the QMAP benchmark.

## Results

### Benchmark construction

To overcome the limitations of the general-purpose PepBenchmark^23^ for antimicrobial peptide regression, we constructed a domain-specific benchmark based on the updated DBAASP database v3^24^. DBAASP provides a substantially larger and more diverse collection of antimicrobial peptides than GRAMPA^10^, the dataset used in PepBenchmark, making it better suited for robust evaluation of AMP regression models.

Dataset construction followed a series of filtering and standardization steps designed to balance biological relevance with practical usability for machine learning. We retained peptides with no terminal modifications as well as those with commonly observed modifications, including N-terminal acetylation and C-terminal amidation. Peptides containing non-canonical amino acids were excluded unless a corresponding SMILES string was available, with the exception of ornithine and 2,4-diaminobutyric acid, two non-canonical residues frequently observed in antimicrobial peptides. In addition, we retained only sequences with either no intrachain bonds or commonly occurring intrachain bonds, such as disulfide bonds and amidation bonds. As a result, the final dataset includes annotations for terminal modifications, D-amino acids, and intrachain bonds, along with SMILES representations for peptides containing non-canonical residues. All entries were formatted into a machine-learning-ready representation.

For antimicrobial activity, we removed potency annotations corresponding to non-bacterial targets and retained only MIC measurements. Consistent with common practice in the field, all MIC values were converted to micromolar (µM) units. When multiple MIC measurements were available for a given peptide–bacterial target pair, outliers were removed using an interquartile range (IQR) criterion and the remaining values were averaged, following the procedure used in PepBenchmark. In cases where annotations were reported at the strain level, measurements were merged at the species level to obtain a consensus MIC value.

For hemolytic activity, we retained all HC50 measurements targeting mammalian erythrocytes. When multiple annotations were available for a peptide, measurements obtained using human erythrocytes were prioritized. Otherwise, the same IQR-based outlier removal and averaging procedure used for MIC were applied to obtain a consensus HC50 value. All HC50 measurements were also converted to micromolar units. Together, these processing steps yield a benchmark comprising two predictive tasks: MIC regression, which directly measures antimicrobial potency, and hemolytic activity (HC50) regression, which captures cytotoxicity toward erythrocytes. Including both tasks enables joint assessment of efficacy and safety, a central consideration in AMP development.

To ensure long-term usability and comparability, we provide predefined test splits as part of the benchmark. This design allows new training data to be incorporated without altering the evaluation protocol or compromising comparability with previously reported results. Because peptide datasets exhibit substantial variability, even models with similar architectures can show markedly different performance depending on the specific test sequences used. To capture this variability, we define five different test splits, and all models are evaluated across each split, enabling systematic quantification of performance variability.

Finally, existing AMP models differ in both input representation and prediction scope. Some methods incorporate terminal modification information, whereas others operate solely on primary sequence, and some are trained on a single bacterial species while others span multiple targets. To enable fair comparison under these heterogeneous assumptions, we evaluate each model on the largest common task supported by all methods under comparison. In practice, this means that models are compared only on predictions for peptide sequences and bacterial targets that are shared across the evaluated approaches. For example, when comparing a model trained to predict MIC values across multiple bacterial species with a model specialized for E. coli only, evaluation is restricted to the E. coli predictions, ensuring that both methods are assessed on an identical and biologically meaningful task.

### Homology-aware split to ensure generalization is evaluated

In protein and peptide modeling, it is essential to prevent homologous sequences from appearing in both the training and test sets, as such overlap can artificially inflate model performance by enabling memorization rather than genuine generalization^21^.

To quantify potential information leakage between training and test data, we adapted a metric originally introduced in the context of natural language processing tasks^25^, which we refer to as the maximum-identity. For each test sample *i*, we compute the global sequence identity^26^ between that sample and every sequence in the training set, and retain the highest value. Formally, letting *T* denote the set of training sequences and Id(·, ·) the global sequence identity, the maximum-identity for test sample *i* is defined as

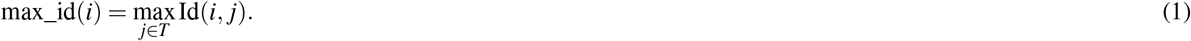

This yields a vector of length *n*_test_, referred to as the maximum-identity distribution, which characterizes the extent of similarity between training and test sequences.

GraphPart^21^ provides a principled mechanism to control sequence leakage through similarity-aware splitting. In this framework, the dataset is represented as a graph in which nodes correspond to sequences and edges connect pairs of sequences whose identity exceeds a predefined threshold. Dataset splitting is then formulated as a graph partitioning problem, ensuring that no training sequence shares an identity above the threshold with any test sequence. This strategy guarantees that the maximum pairwise train–test identity is bounded by the chosen threshold. The impact of this approach, relative to commonly used random train–test splits, can be illustrated by examining the resulting maximum-identity distributions (Fig 1). Random splits yield distributions that are strongly shifted toward higher identities and exhibit substantial upper tails, whereas homology-based splits produce sharply truncated distributions whose upper tails do not extend beyond the threshold, thereby enforcing strict control over train–test similarity and preventing inadvertent information leakage.

**Figure 1.**
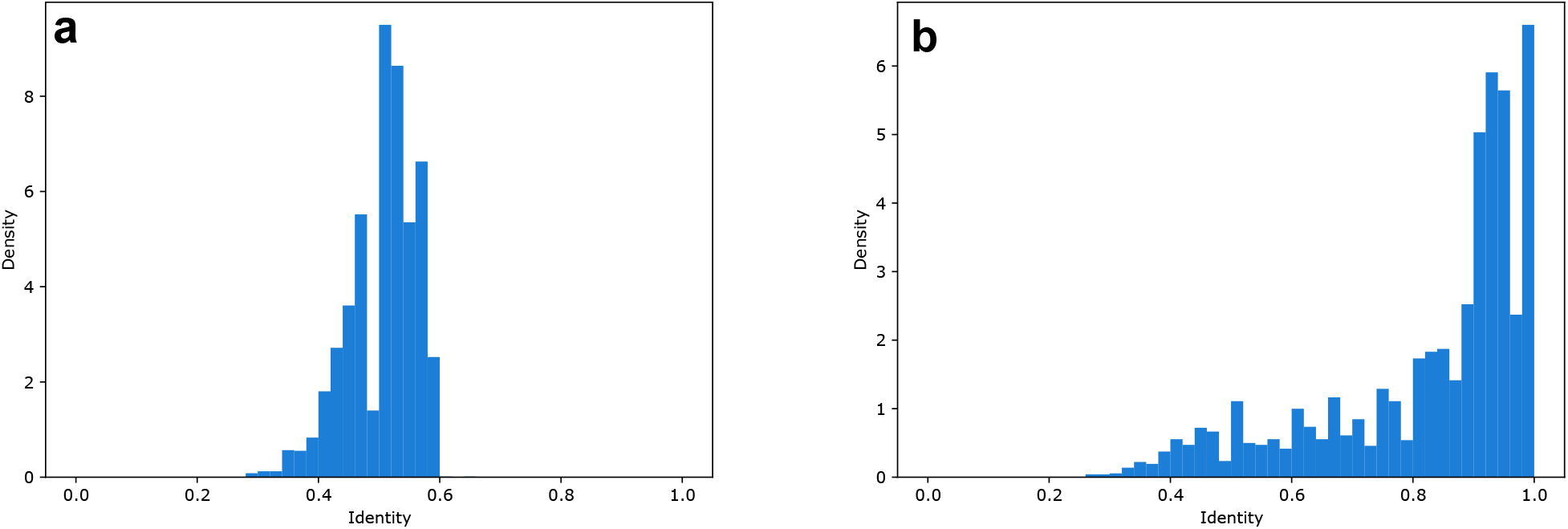
Maximum-identity distributions between the test and training sets on the DBAASP dataset under two split strategies, shown as probability density histograms. **a**, Homology-aware split with a threshold of 60%. The distribution is sharply truncated at the threshold, confirming strict control over train–test sequence similarity. Median: 50.0%; 90th percentile: 57.1%. **b**, Standard random split. The distribution is strongly shifted toward high identity values, with a pronounced spike near 1.0 attributable to sequences in DBAASP that differ only by N- or C-terminal modifications and are therefore identical at the sequence level. Median: 90.0%; 90th percentile: 100.0%.

A key design goal of our benchmark is to enable evaluation of models trained on arbitrary external datasets, which may contain sequences highly similar to those in the benchmark test sets and thus violate strict train–test homology minimal distance. To enforce the similarity threshold in this setting, we compute pairwise sequence identities between the external training data and the benchmark test sequences, and remove any training samples whose identity to a test sequence exceeds the specified threshold.

Applying GraphPart directly to the DBAASP dataset revealed a practical limitation: the dataset’s dense similarity structure causes strict graph partitioning to yield an empty test set, leaving no sequences for meaningful evaluation. To address this issue, we modified the splitting procedure by (i) applying a community detection algorithm to group sequences based on similarity and (ii) randomly selecting entire clusters to construct the test set, whereas GraphPart performs this step deterministically. Similarly to GraphPart, a final filtering step removes training samples whose similarity to any test sample exceeds the specified threshold. This procedure is schematized in Fig 2. Using this strategy with different random seeds, we generated five distinct similarity-aware test sets, which together constitute the QMAP benchmark splits. This similarity-aware splitting approach aims to preserve most training sequences while ensuring a realistic evaluation of generalization, which is particularly important given the limited size of annotated AMP datasets.

**Figure 2.**
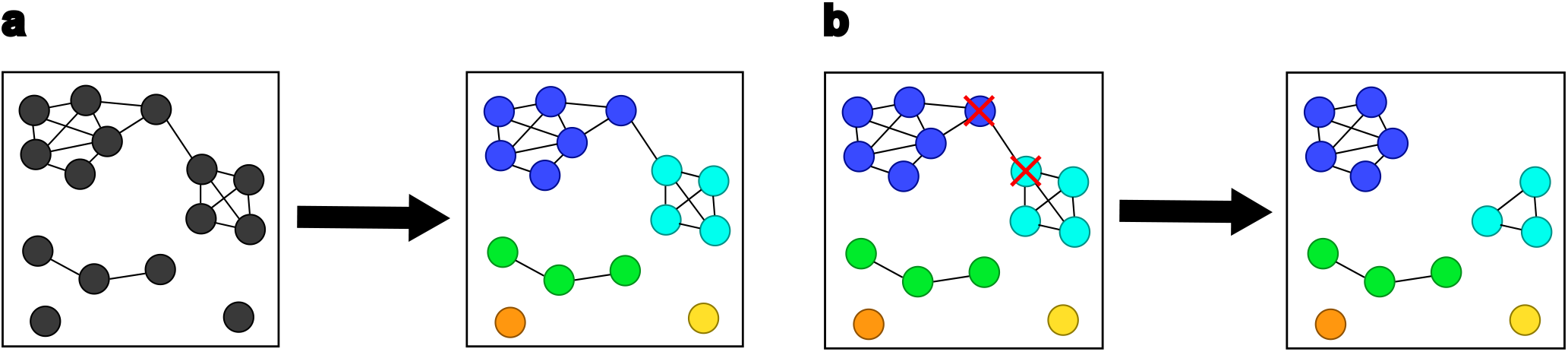
Illustration of the homology-based split algorithm following graph construction, where nodes represent sequences and edges connect pairs sharing identity above the threshold. **a:** Community detection: the Leiden algorithm with the Modularity objective is applied to the graph, partitioning sequences into communities, each denoted by a distinct colour. **b:** Node filtering: communities are randomly assigned to the training or test set. In this example, blue and cyan communities are assigned to different sets. Sequences sharing an edge across the train–test boundary (marked with a red cross) are removed to ensure a strict separation with respect to the identity threshold.

### Threshold identification

The identity threshold introduced in our framework acts as a hyperparameter, and its choice is therefore non-trivial. Setting the threshold too high results in overly optimistic evaluation performance, as test samples remain closely related to training samples. Conversely, setting the threshold too low leads to overly conservative estimates by unnecessarily restricting the diversity of the training set: fewer sequence clusters are observed during training, which can in turn degrade real-world performance (Fig 3a). Importantly, choosing a relation threshold is equivalent to specifying the expected distance between training data and deployment data. This mirrors established practices in protein bioinformatics, such as the choice of a BLOSUM substitution matrix^27^. Whereas a BLOSUM matrix implicitly encodes assumptions about evolutionary distances, the relation threshold explicitly encodes assumptions about deployment distance. In this sense, threshold selection is not arbitrary but reflects prior beliefs about the conditions under which a model will be applied.

**Figure 3.**
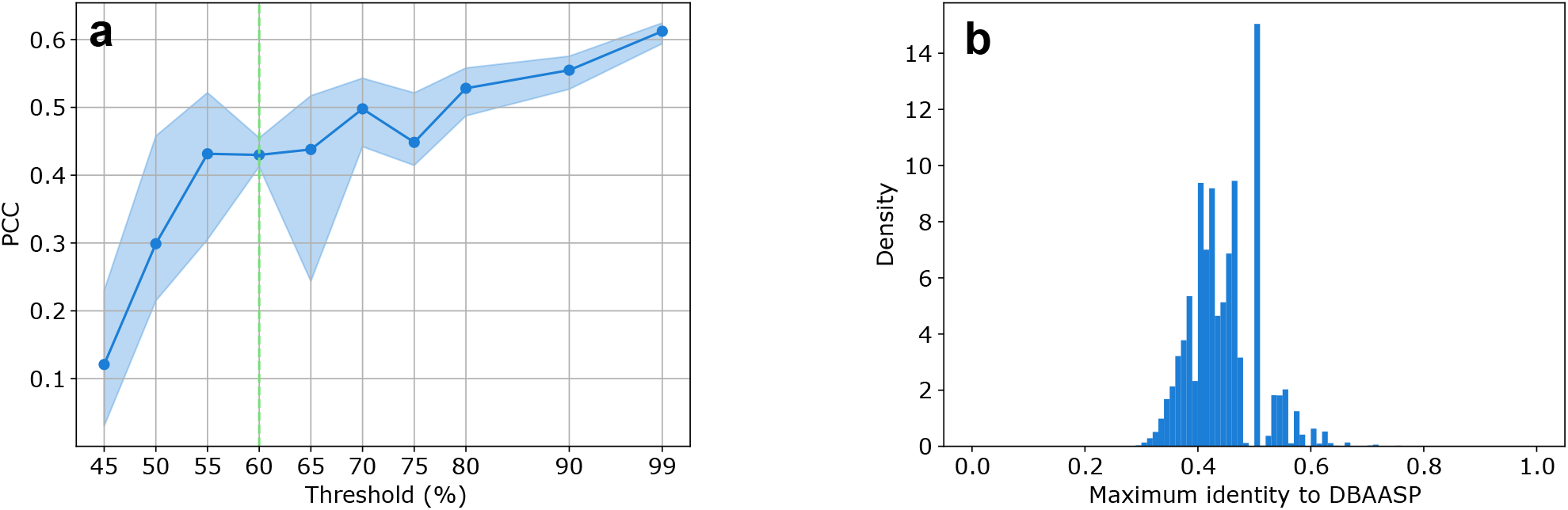
Threshold selection for the homology-based split. **a:** Pearson correlation coefficient (PCC) of a linear probing model trained on ESM2-650M embeddings, evaluated across increasing identity thresholds splits. The DBAASP dataset is partitioned using the homology-based split strategy at each threshold, where a higher threshold permits greater sequence similarity between the training and test subsets, resulting in a less stringent separation. The experiment is repeated five times per threshold with different random splits, and the shaded area represents the standard deviation across runs. PCC increases consistently with the threshold, indicating that less stringent splits inflate apparent model performance. The green dashed vertical line marks the selected threshold of 60%, used to construct the benchmark. **b:** Probability density histogram of the maximum sequence identity between the Peptide Atlas database^28^ and the DBAASP dataset. The 99th percentile of this distribution is 61.5%, indicating that 99% of known peptides share at most 61.5% identity with DBAASP, motivating the selection of the 60% threshold as a biologically grounded and conservative separation boundary.

Models developed to predict MIC or HC50 are commonly used either to screen large, heterogeneous peptide libraries or as reward functions within generative design pipelines. In both settings, inference-time sequences originate from diverse sources and are typically far removed from the relatively small, curated datasets used for training. As a result, the expected distance between training and deployment sequences is large, indicating that evaluation should be conducted under a relatively low identity threshold.

To operationalize this intuition, we estimate the expected maximum-identity distribution between deployment sequences and the training dataset. We use the largest available natural peptide collection, Peptide Atlas^28^, as a proxy for the space of naturally occurring peptide sequences. We then compute the maximum identity between peptides in this dataset and those in our antimicrobial peptide dataset (Fig 3b).

We select an identity threshold such that the maximum sequence identity between training and test samples does not exceed the similarity expected between the training set and realistically distant sequences. To estimate this upper bound, we use the 99th percentile of the maximum-identity distribution between PeptideAtlas and DBAASP, providing a conservative and noise-robust criterion. As the 99th percentile of this distribution is 61%, we adopt a slightly more conservative threshold of 60% sequence identity for all benchmark splits.

### Evaluation of MIC regression progress under controlled benchmarking

To assess progress in AMP MIC regression under a controlled and reproducible evaluation framework, we reviewed existing approaches and identified a limited subset of models for which code and trained configurations were publicly available. Among these, we successfully reproduced the reported results of two representative methods: J. Witten et al.^10^ and BERT-AmPEP60^17^, both of which use a neural network to predict the antimicrobial activity of peptides against *E. coli*. To provide an additional point of reference, we implemented a simple baseline consisting of a linear probe trained on embeddings from ESM-650M, a pretrained protein language model (PLM)^29^.

We computed a set of complementary metrics to obtain a holistic view of predictive performance. Correlation-based metrics were used to evaluate how well models preserve relative activity ordering:

- *R*^2^, the coefficient of determination, measures the proportion of variance in observed MIC values explained by the model predictions.
- Kendall’s *τ*, a rank-based correlation metric, quantifies concordance between predicted and observed activity rankings and is robust to outliers and non-linear relationships.
- Spearman’s correlation captures the monotonic relationship between predicted and observed MIC values based on rank ordering, without assuming linearity.
- Pearson’s correlation quantifies linear correlation between predicted and observed MIC values, emphasizing global linear agreement.

We also report error-based metrics to assess absolute predictive accuracy:

- RMSE penalizes large prediction errors more strongly and reflects overall prediction precision.
- MAE provides an interpretable measure of the average absolute deviation between predicted and observed MIC values.

Higher values of correlation-based metrics indicate better preservation of relative activity ordering, whereas lower values of error-based metrics indicate more accurate quantitative predictions.

When evaluated on the QMAP benchmark, both deep learning approaches outperform the linear baseline across all metrics. However, the performance gap between the two neural models is modest (Fig 4a,b). In addition, the performances obtained on QMAP differ substantially from those reported in the original studies. To assess the impact of evaluation protocols, we compared model performance on QMAP with performance obtained using a standard random split strategy (Fig S1a). Models evaluated on random splits achieved substantially higher performance across all metrics. One possible explanation is that random partitioning preserves strong sequence similarities between training and test samples, enabling models to rely on local interpolation between homologous peptides rather than learning relationships that generalize beyond syntactic similarity.

**Figure 4.**
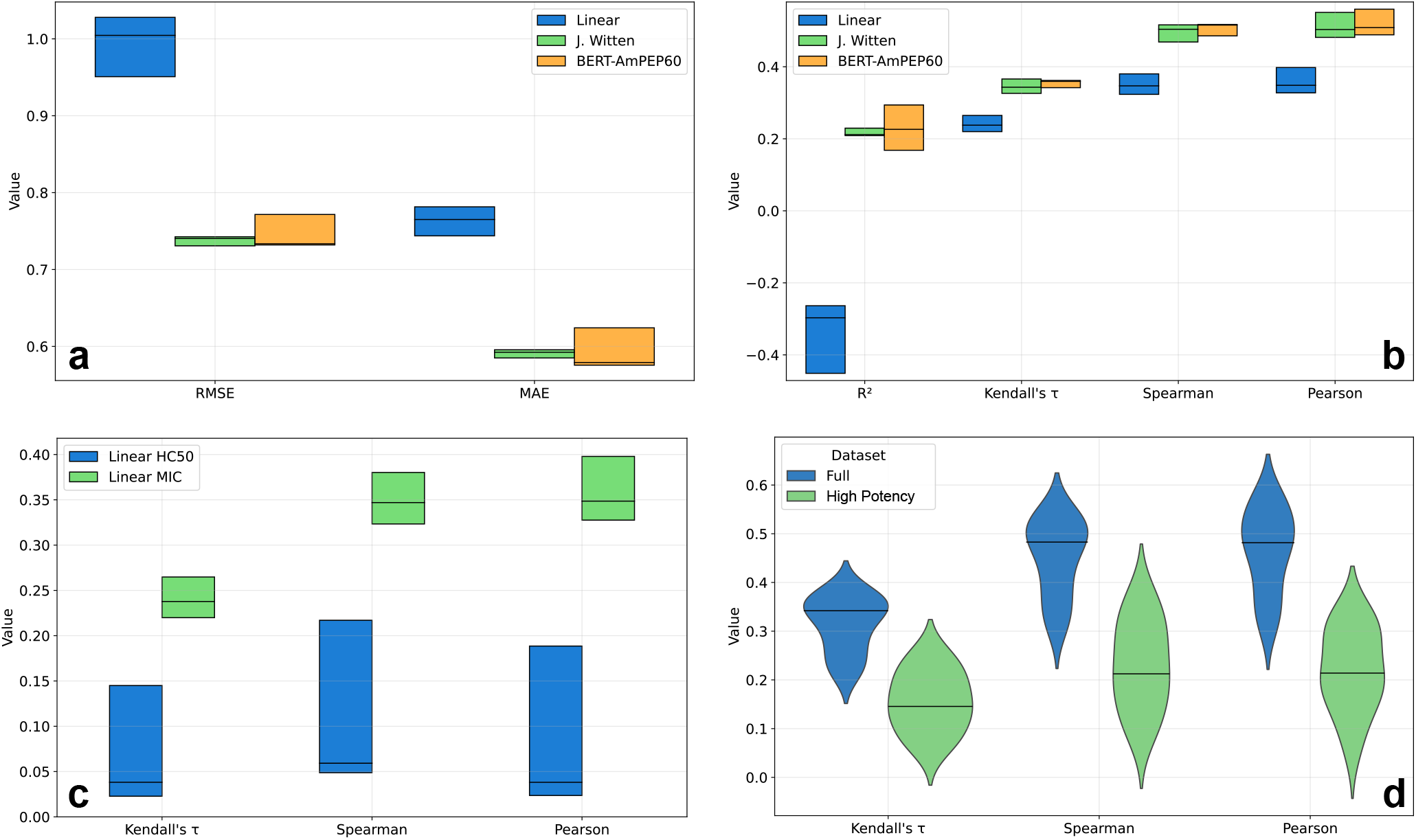
Benchmark performances of the linear baseline, J. Witten et al.^10^, and BERT-AmPEP60^17^. Unless otherwise stated, all panels aggregate results across the five benchmark splits, and each box represents a custom plot where the top and bottom edges correspond to the second highest and second-lowest values respectively, and the horizontal line denotes the median. Linear refers to the linear probing model trained on ESM2 650M embeddings. **a:** Loss metrics (lower is better). **b:** Correlation metrics (higher is better). **c:** Correlation metrics for the linear baseline on the hemolytic regression task (HC50) compared to the same model trained and evaluated on the MIC task, across the same five benchmark splits. The *R*^2^ score is omitted as strongly negative values distorted the axis. **d:** Violin plots comparing correlation performances of the three models (Linear, J. Witten et al., and BERT-AmPEP60) on the full dataset versus the high-potency subset (MIC *<* 10 *µ*M), aggregating results across all three models and five benchmark test sets (15 points per violin). The horizontal line denotes the median. The *R*^2^ metric is omitted for visual clarity as values were strongly negative.

To further investigate this possibility, we analyzed the relationship between prediction error and the distance separating each test peptide from its nearest training sample (Fig S1b). Across models, prediction error increased with increasing distance to the training set, indicating that peptides located closer to previously observed sequences were easier to predict. These observations suggest that random split strategies may inflate performance estimates by favoring similarity-driven prediction shortcuts. In contrast, homology-aware partitioning reduces this effect by enforcing greater separation between training and evaluation samples, thereby providing a more stringent assessment of model behavior. Without standardized evaluation splits and similarity-aware partitioning strategies, the extent to which reported performances depend on local sequence similarity remains difficult to quantify and has likely been underestimated in prior studies.

Taken together, these results suggest that progress in AMP MIC regression over the past six years has been more limited than previously perceived. Inconsistent evaluation protocols and split-dependent performance estimates have likely obscured this stagnation, emphasizing the importance of standardized benchmarking and controlled, homology-aware test sets for accurately assessing advances in the field.

### Regime-specific limitations in MIC prediction

Beyond aggregate performance metrics, a key advantage of the QMAP benchmark is its ability to reveal regime-specific failure modes that are directly relevant to real-world AMP discovery. In practice, the utility of a model is often determined not by average performance, but by its behavior in high-impact regions of the task space.

To illustrate this phenomenon, we evaluated model performance in a high-potency regime by restricting the test sets to low-MIC peptides. Low-MIC peptides are defined as those with a MIC annotation below 10 µM for at least one target, corresponding to approximately half of the training and benchmark datasets. Peptides with a MIC below 10 µM are considered hits in some drug discovery pipelines^30^ — the primary candidates for further experimental validation and therapeutic development. Despite reasonable aggregate performance across the full dataset, we observed a pronounced degradation in predictive accuracy within this high-potency subset (Fig 4d). This limitation is particularly concerning because highly potent peptides are the primary focus of *in silico* screening and generative design pipelines.

To further investigate this phenomenon, we performed two stratified analyses across MIC-defined potency regimes. First, we evaluated model performance separately on high-potency (HP; MIC *<* 10 µM), intermediate-potency (IP; 10 ≤ MIC *<* 100 µM), and low-potency (LP; MIC ≥ 100 µM) subsets of the benchmark datasets (Fig S2a). Across all evaluated models, predictive performance progressively deteriorated with increasing peptide potency, with the HP subset consistently exhibiting the lowest correlations. Notably, this trend was not explained by differences in sample availability, as the HP subset contained more samples than both the IP and LP subsets, yet remained the most challenging regime to predict. To determine whether this limitation could be mitigated through regime-specific enrichment, we retrained and evaluated models exclusively within each potency subset (Fig S2b). Although overall performance decreased due to the reduced training set sizes, the same monotonic trend persisted, with predictive accuracy remaining the lowest in the HP regime.

The persistence of this effect across both aggregate and regime-specific training strategies suggests that the reduced predictability of highly potent peptides is not solely attributable to conventional learning dynamics associated with sample count or target-range specialization. One possible explanation is the presence of dataset-level biases or experimental heterogeneity within DBAASP, where differences in assay protocols, laboratory conditions, or reporting practices may introduce substantial variability in measured MIC values. In addition, the currently available AMP datasets may incompletely sample the underlying sequence and activity landscape, particularly within highly potent regions of sequence space. Alternatively, the observed trend may reflect an intrinsic biological property of antimicrobial peptides, whereby highly potent peptides exhibit a more heterogeneous or locally sensitive sequence–activity relationship. Under such a scenario, small sequence variations could induce disproportionately large changes in antimicrobial activity, resulting in a less smooth mapping between sequence and potency. These explanations are not mutually exclusive, and further investigation will be required to determine the relative contributions of experimental, dataset, and biological factors underlying this phenomenon.

These results highlight a systematic limitation shared across current modeling approaches and underscore the importance of evaluating models under task-relevant operating conditions. They further motivate the use of environment-specific evaluations to uncover vulnerabilities that remain invisible under aggregate metrics alone.

### Challenges in hemolytic activity prediction

Hemolytic activity is evaluated as a regression task within the benchmark, using HC50 values. For consistency with MIC regression experiments, we adopt the same predefined test splits and independence constraints, enabling direct comparison across tasks.

To establish a reference point for this task, we evaluate a baseline model consisting of a linear probe trained on fixed embeddings from ESM-650M. While we sought to include existing methods for comparison, no prior work on hemolytic activity regression provided reproducible code, limiting our evaluation to this standardized baseline.

As shown in Fig 4c, predictive performance on hemolytic activity for the linear baseline is substantially lower than that observed for MIC prediction across all evaluation metrics. This performance gap highlights the intrinsic difficulty of HC50 regression relative to potency prediction.

We hypothesize that this challenge arises from several factors, including the smaller dataset size (800 peptides, compared to 4000 for E. coli MIC) and greater heterogeneity in experimental conditions and peptide mechanisms. Together, these factors likely constrain the amount of learnable signal available to supervised models.

Furthermore, the limited performance of the linear probe suggests that current pretrained protein embeddings capture only weak linearly accessible information relevant to hemolytic activity. Taken together, these results establish a baseline for HC50 prediction on the benchmark and identify hemolytic regression as an open and underexplored challenge for future AMP modeling efforts.

## Discussion

Despite significant advances in AMP prediction, quantitative comparison between models has remained challenging due to differences in dataset construction, split strategies, and evaluation protocols.

Our benchmark addresses these limitations by providing a predefined, homology-aware evaluation framework composed of fixed testing data subsets and a procedure to ensure strict minimal distance between training and test sets to ensure generalization is evaluated. This approach guarantees consistency across models and training datasets and allows direct, reproducible comparison of MIC and hemolytic regression performance. To capture performance variability, multiple independent evaluation sets are included. Furthermore, the benchmark incorporates annotations for terminal modifications, intrachain bonds, D-amino acids, and SMILES strings for non-canonical amino acid containing peptides, ensuring compatibility with current and emerging AMP modeling approaches. Future extensions of QMAP could incorporate domain-combined and chimeric peptide architectures, which are increasingly relevant in modern AMP research^31^, provided that dedicated databases with MIC and HC50 annotations for such constructs become available.

Evaluation of existing MIC prediction methods for *E. coli* showed that apparent improvements over the past several years are more modest than previously reported. This outcome reflects the influence of dataset split strategies and the lack of standardized benchmarks, which can inflate perceived generalization performance. These results suggest that a considerable fraction of the reported performance gains may arise from similarity-driven interpolation enabled by conventional random split strategies. The challenge of generalization is even more pronounced for hemolytic activity regression, where smaller training datasets limit predictive performance and reduce the capacity to learn robust features.

An important limitation of this finding is that only two reproducible MIC prediction models could be systematically evaluated on QMAP, spanning approximately six years of methodological development. Although additional approaches have been reported in the literature, most lacked publicly available and functional code for reliable reproduction. The observed trend should therefore be interpreted cautiously, as it may not fully capture the breadth of progress across the field. These observations further emphasize the importance of standardized benchmarks, publicly accessible implementations, and reproducible evaluation pipelines for enabling transparent and comparable assessment of future AMP prediction methods.

Across all evaluated methods, our benchmark revealed a consistent limitation: predictive performance deteriorates sharply for highly potent peptides, a critical regime for *in silico* screening and generative design applications. This finding emphasizes an important direction for future research aimed at improving the discovery of effective antimicrobial peptides. In addition, the linear probing baseline trained on embeddings from a foundational protein language model exhibited limited performance, suggesting that these general embeddings only weakly capture information related to true hemolytic activity.

The limited size of available datasets, particularly for HC50 regression, indicates that predictive performance could benefit from richer, more representative features derived from pretrained foundational models or other unsupervised learning approaches. However, the weak correlation observed between protein language model embeddings and hemolytic activity highlights a key challenge: features learned from general protein sequences do not linearly transfer to bioactive peptides. This suggests a need for task-specific or peptide-adapted representations better suited to bioactive peptides. The team behind PepBenchmark found that continuing the masked language model pretraining on bioactive peptide datasets of foundational models can improve performance in certain scenarios, mainly for classification tasks^23^. Drawing from analogous advances in computer vision, foundation models have demonstrated that well-designed pretraining schemes can yield linearly separable representations, enabling greater sample efficiency on downstream tasks^32^ — a property that would be particularly valuable given the limited size of AMP datasets. This motivates the development of pretraining schemes tailored to bioactive peptides, with the goal of producing representations that more directly encode the properties relevant to antimicrobial and hemolytic activity.

Ultimately, QMAP does not itself provide new peptide candidates or mechanistic insights, but offers a standardized foundation that facilitates and accelerates such downstream work.

## Methods

### Building the benchmark

We downloaded the DBAASP database using its official API and applied a series of filtering and normalization steps to construct a machine-learning-ready benchmark. We first removed peptides reported as multimers or multi-peptide complexes, retaining only monomeric peptides. Peptides with N-terminal modifications other than acetylation or C-terminal modifications other than amidation were excluded.

Non-canonical amino acids were handled as follows. Ornithine residues, originally represented as X, were replaced by O, and 2,4-diaminobutyric acid (DAB) residues were replaced by B. When present in their D-stereoisomeric form, these residues were represented as o and b, respectively. Any remaining non-canonical amino acids led to sequence removal unless a corresponding SMILES string was available. Finally, peptides containing intrachain bonds other than disulfide or amide bonds were excluded. After these filtering steps, the dataset contained 18,033 peptide sequences.

We retained only bacterial minimum inhibitory concentration (MIC) measurements. Whether a target corresponded to a bacterial species was determined using the ete4 package^33^ with NCBITaxa. All MIC values were converted to micromolar (µM) units. When annotations were provided at the strain level, strain information was discarded to obtain species-level annotations. Many MIC values were reported as ranges or semi-defined bounds (e.g., “less than” or “greater than”). Undefined lower and upper bounds were represented as 0 and ∞, respectively. To derive a consensus MIC value, all definite bounds were treated as measurements, outliers were removed using an interquartile range (IQR) criterion, and values outside [Q1 - 1.5× IQR, Q3 + 1.5 × IQR] were excluded. The remaining values were then averaged. Since variability in reported MIC values primarily reflects differences in experimental conditions and bacterial isolates rather than measurement uncertainty, aggregating multiple annotations via the mean provides a robust estimate of central tendency that preserves the correlation with peptide efficacy necessary for predictive modeling.

An identical filtering and averaging procedure was applied to hemolytic activity (HC50) measurements. However, when a human erythrocyte measurement was available, it was directly assigned as the consensus value without averaging. Otherwise, a consensus HC50 value was computed by averaging measurements obtained from mammalian erythrocytes after IQR-based outlier removal.

This process yields the DBAASP-derived dataset used throughout this study and for benchmark construction. The resulting dataset includes MIC annotations for 492 bacterial species, hemolytic activity measurements, annotations for N- and C-terminal modifications, intrachain bond information, and SMILES strings for peptides containing non-canonical amino acids.

Benchmark splits were constructed using our similarity-aware, graph-based splitting procedure. For each split, a train–test partition was generated, but only the test sets were retained, as the corresponding training sets can be trivially recovered by removing test set peptides and their similar sequences from the full DBAASP dataset using the provided toolkit. Five such splits were produced using different random seeds (1, 3, 7, 12, and 404). The collection of these five test sets constitutes the QMAP benchmark. Both the formatted DBAASP dataset and the predefined benchmark splits were made publicly available via the Hugging Face platform.

### Using the benchmark

QMAP is designed to be training-set agnostic: any dataset can be used for training provided that the minimum homology distance constraint between training and test sets is satisfied. For each predefined test split, the provided toolkit identifies all sequences in a given training dataset that are similar to test set peptides, allowing them to be safely removed prior to training. This ensures that no sequence leakage occurs between training and test sets. As all models are evaluated on identical test sets with the same homology constraint enforced, results obtained under different training configurations remain directly comparable.

### Python package accelerated with Rust engine

To facilitate the adoption of the benchmark, we released a Python package distributed via the `pip` package manager. The package includes the formatted DBAASP dataset used in this study, along with the five predefined benchmark test splits. It also provides a suite of utility functions to streamline benchmark usage, including automatic computation of evaluation metrics and generation of masks identifying training samples that satisfy the benchmark’s maximal similarity constraints for each split.

In addition, the package includes a toolkit for peptide-specific data manipulation. This toolkit provides functionality analogous to commonly used utilities in machine learning workflows, but tailored for peptide data. Such functionalities include a train–test splitting function similar to that in `scikit-learn`^34^, as well as functions for constructing sequence similarity graphs and identifying clusters within these graphs.

Several of the required operations are computationally intensive, particularly those involving large-scale sequence similarity calculations. To address this, we developed an optimized pairwise identity computation engine implemented in Rust, referred to as the *pwiden* engine. This engine enables efficient computation of full pairwise identity matrices, construction of identity graphs, generation of training-set masks that enforce the maximal similarity thresholds with respect to the test set, and more. It leverages SIMD operation thanks to `Parasail`^35^, an efficient C library to align sequences, and multi-thread parallelism thanks to the `rayon` library, available as a Rust *crate*. All of these capabilities are exposed through Python bindings and integrated seamlessly into the released package.

### Data leakage quantification

The function and biological activity of a peptide are largely determined by its amino acid sequence, which in turn defines its structure and physicochemical properties. As a consequence, peptides with similar sequences are often closely related evolutionarily and are therefore likely to exhibit similar functional behavior. Evaluating a model on a test set that contains sequences closely related to those seen during training is thus expected to yield artificially inflated performance compared to evaluation on more distant sequences.

To quantify this effect, we measure the degree of homology between training and test datasets using the maximum-identity metric. We then compare the resulting maximum-identity distributions obtained from two different splitting strategies: our adaptation of GraphPart, which explicitly enforces homology-aware separation, and a random train–test split, which serves as a naive baseline. For both strategies, we use the same underlying dataset as that employed for constructing the benchmark in order to generate the maximum-identity distribution figures.

For the graph-based splitting strategy, we used the default parameters provided in the `qmap-benchmark` package: a sequence identity threshold of 60%, a test set ratio of 20%, post-filtering enabled, and the same alignment parameters as those used in the definition of the maximum-identity metric. For the random split baseline, we randomly selected 20% of the dataset as the test set, without imposing any similarity constraints.

#### Maximum identity metric

For each test sequence, we compute its sequence identity with every sequence in the corresponding training set and retain only the highest value. This results in a set of scalar values, one per test sequence, which together form the maximum-identity distribution. Pseudocode for computing this metric is provided in Listing 1. We visualize this distribution using a density histogram and report aggregate statistics, including the median and the 90th percentile.

The interpretation of this metric is straightforward: distributions shifted toward higher maximum identities indicate a greater degree of similarity between training and test sets and, consequently, a higher risk of information leakage.

Sequence alignment and global identity computation are performed using the Needleman–Wunsch algorithm^26^. We parameterize the alignment with the BLOSUM45 substitution matrix, an open gap penalty of −5, and a gap extension penalty of −1.

### Dataset Partition algorithm

The first step in constructing the data split is to build a sequence similarity graph, in which each peptide sequence is represented as a node, and an edge is introduced between two nodes if their pairwise sequence identity exceeds a user-defined threshold. This requires the computation of all pairwise sequence identities. The procedure is formalized through the adjacency matrix

**Listing 1.**
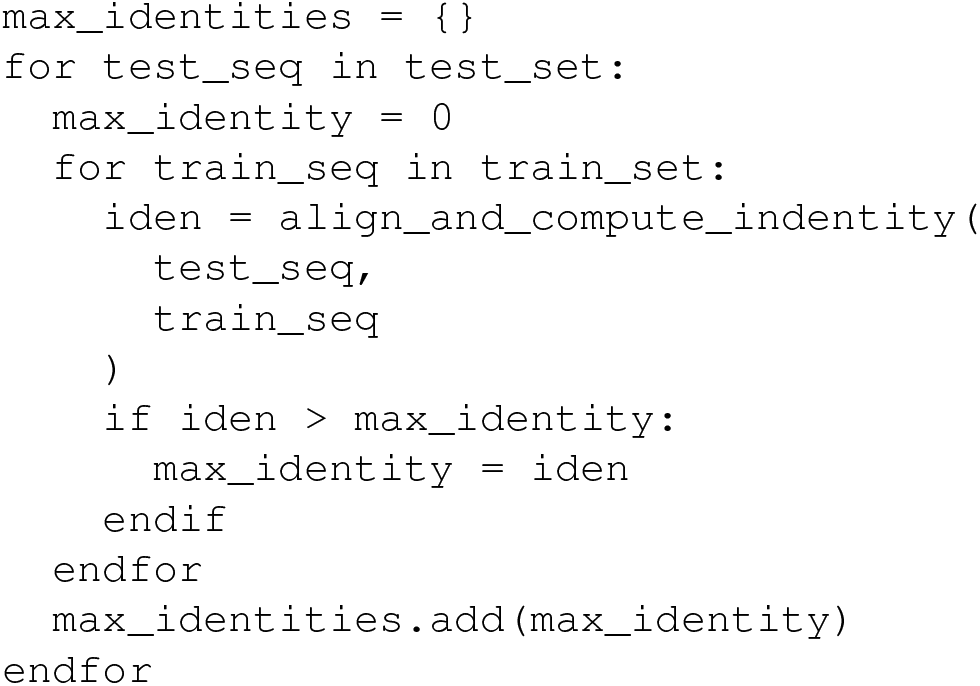
Pseudocode for computing the maximal identity metric. For each test sequence, the identity to every training sequence is computed via global pairwise alignment, and the maximum value is retained, yielding the identity to the most similar sequence in the training set.

defined in (2) for a given threshold *t*, where the adjacency entry *A*_*i, j*_ indicates whether the identity between sequences *i* and *j* exceeds *t*. This forms the adjacency matrix *A* ∈ ℝ^*N*×*N*^ where *N* is the number of sequences in the dataset.

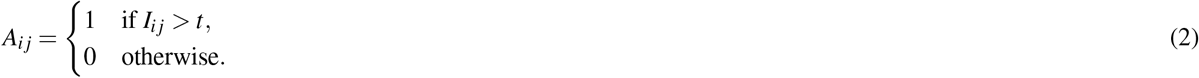

To make this step computationally tractable, pairwise identity computation is accelerated using a compiled Rust implementation. Exploiting the symmetry of the identity matrix, only half of the pairwise comparisons are evaluated, and only the resulting edges are stored in memory, substantially reducing computational and memory overhead.

Communities in the similarity graph are then identified using the Leiden algorithm^36^, which guarantees that all detected communities are internally connected. Importantly, this algorithm does not require hyperparameter tuning, thereby limiting potential sources of bias introduced by arbitrary parameter choices during dataset partitioning. We use the `leidenalg` package, which provides a high-performance C++ backend with a convenient Python interface, and adopt the modularity-based vertex partition for community detection.

The dataset subsets are constructed at the community level using a greedy procedure. The test subset is assembled first by uniformly sampling and aggregating communities until the desired test set size is reached. The training subset is then formed by aggregating the remaining communities.

Finally, a post-filtering step is applied to strictly enforce the similarity constraint. Each training sequence is aligned against all test sequences, and any training sequence that exceeds the predefined identity threshold with at least one test sequence is removed.

### Ideal threshold

To guide the selection of an appropriate sequence identity threshold, we constructed a large reference distribution of sequence similarities. The PeptideAtlas dataset was assembled by downloading all the latest builds from the PeptideAtlas database. The DBAASP dataset distributed with the package served as the reference set. Because PeptideAtlas is approximately 200 times larger than DBAASP, this comparison provides a realistic estimate of the similarity distribution between curated AMP datasets and large, general peptide repositories. Maximum identities between the two datasets were computed using our optimized `pwiden` engine. The resulting distribution was visualized as a histogram using `matplotlib`, and the 99th percentile was computed using the `numpy` package.

To assess the impact of the identity threshold on model evaluation, we performed a series of experiments across a range of thresholds. For each threshold, the DBAASP dataset was partitioned into training and test sets using our graph-based splitting method. Sequence embeddings were generated using ESM2-650M, and MIC labels for E. coli were sampled from the DBAASP dataset distributed with the package. A linear regression model without regularization was trained on each resulting training set, and performance was evaluated on the corresponding test set using the Pearson correlation coefficient (PCC). Each experiment was repeated five times per threshold to quantify performance variability.

### Evaluation of related works

For each external method evaluated, we first verified that the originally reported results could be faithfully reproduced. Only minimal modifications were made to the released source code to enable evaluation on the QMAP benchmark. In practice, this involved installing the qmap-benchmark package and iterating over the five predefined benchmark test splits. For each split, the original training dataset was subsampled to enforce a minimum sequence identity distance of 60% with respect to the selected benchmark test set. This filtering step is fully automated within the package and requires only a few lines of code. Apart from this constraint, the original training procedures of the evaluated methods were preserved without modification. Model predictions were then generated on the benchmark test sets and compared to the corresponding ground-truth labels using the metric suite provided by the QMAP-benchmark package.

In addition to previously published methods, we implemented a linear baseline model for comparison. Each peptide sequence was encoded using embeddings from the ESM2-650M protein language model. A linear regression model without regularization was trained using the scikit-learn package to predict base-10 logarithm–transformed MIC values. The training data consisted of the featurized DBAASP dataset distributed with the benchmark package. As with the external methods, training sequences exceeding the allowed similarity threshold with the test set were removed prior to training.

Results are presented using a compact box-style visualization designed to improve readability by reducing the influence of extreme values. For each model and metric, the upper and lower bounds of the box correspond to the second-highest and second-lowest values, respectively, while the central line denotes the median performance across the five benchmark splits.

To assess model performance in the high-potency regime, we evaluated the same regression approaches on a high-potency subset of the benchmark. This subset was defined by selecting peptides with at least one MIC measurement below 10 µM. All other aspects of the experimental protocol were left unchanged, and the training data remained identical to those used for the full benchmark. To facilitate comparison between full-benchmark and high-potency evaluations, results were summarized using violin plots aggregating performance across all three models and the five benchmark splits, yielding a total of 15 measurements per condition.

To further characterize the relationship between peptide potency and predictive performance, we conducted a stratified analysis across three potency levels: high-potency (HP; MIC *<* 10 µM), intermediate-potency (IP; 10 ≤ MIC *<* 100 µM), and low-potency (LP; MIC ≥ 100 µM). For each stratum, model performance was evaluated by restricting the test sets to peptides falling within the corresponding MIC range, while the training data remained identical to those used for the full benchmark evaluation. In a complementary analysis, models were also retrained on data restricted to each potency stratum, with the test sets restricted accordingly, to assess whether training on potency-matched data alters the observed performance trends. Results were summarized using violin plots aggregating performance across all three models and the five benchmark splits.

## Data availability

Dataset available on Hugging Face: https://huggingface.co/datasets/anthol42/qmap_benchmark_2025

Package installable using the pip tool: `pip install qmap-benchmark`

Code for the package, or reproduce figures is available on GitHub: https://github.com/anthol42/QMAP

## Acknowledgments

JC acknowledges the support of the Canadian Biomanufacturing Research Fund (CBRF) and, specifically, the PandemicStopAI project: an accelerated response to bacterial pandemics. He also acknowledged the funds for the CQDM (Quebec Consortium for Drug Discovery). PG is supported by by the NSERC Discovery grant RGPIN-2020-07223. AL acknowledges funding from the Natural Sciences and Engineering Research Council of Canada (NSERC) through a Canada Graduate Scholarship - Master’s (CGS M).

## Supplementary Material

**Figure S1.**
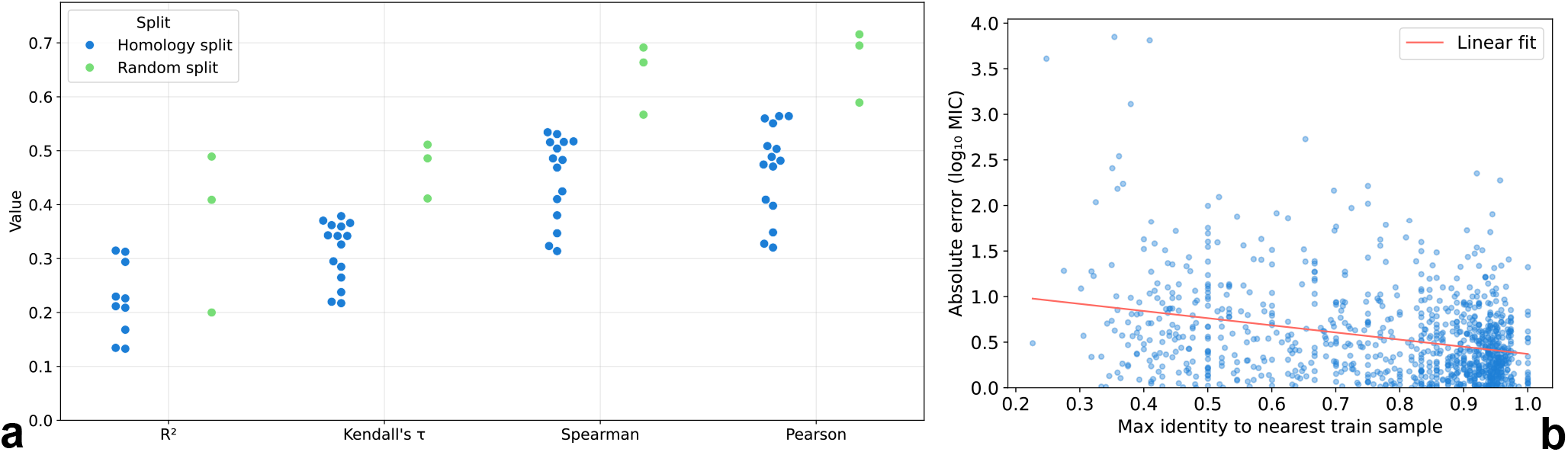
Visualization of potential overfitting when no homology-based split is applied. **a**: Swarm plot of correlation metrics (*R*^2^, Kendall’s *τ*, Spearman, and Pearson) for models trained and evaluated using either a homology split or a random split. For the homology split, the three evaluated models are each assessed across five benchmark sets, yielding 15 individual dots. For the random split, each of the three models is evaluated on a single randomly obtained test set, yielding 3 individual dots. **b**: Scatter plot of the absolute prediction error (*log*_10_ MIC) as a function of the maximum sequence identity to the nearest training sample, for the linear model trained on the random split. A linear regression fit (red line) reveals a descending trend (Pearson r = -0.31), indicating that predictions deteriorate for samples with low identity to the training set.

**Figure S2.**
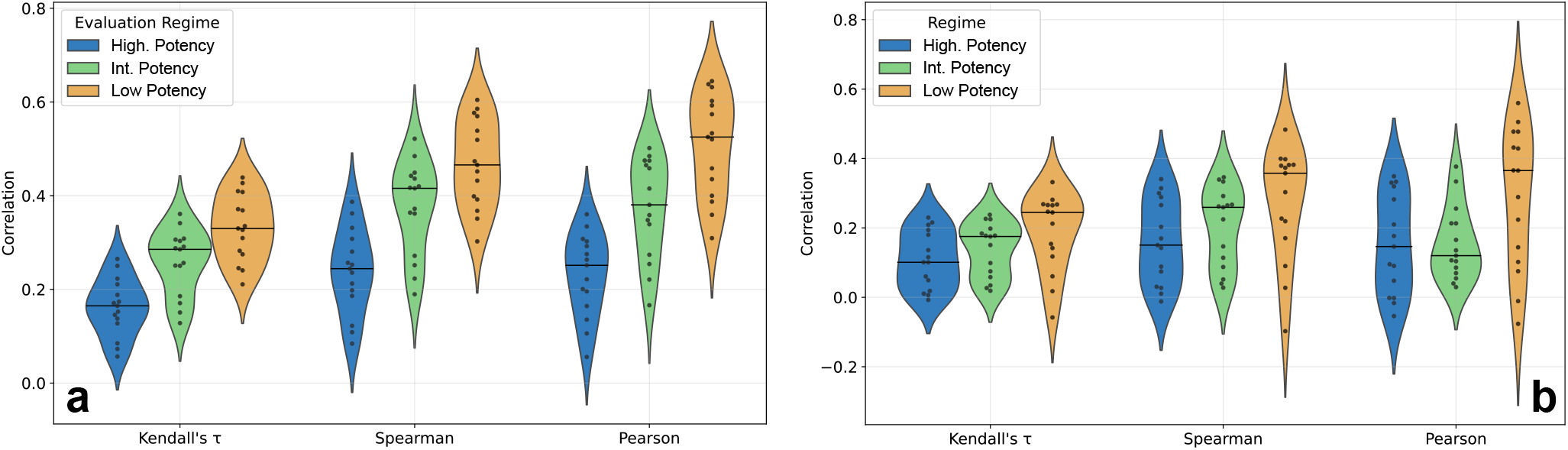
Investigating the high-potency regime collapse. **a:** Violin plots with overlaid swarm plots representing stratified correlation scores (Kendall’s *τ*, Spearman, and Pearson) for all three models evaluated under the homology split, trained on the full dataset. Each dot represents the correlation achieved on the stratified subset of one of the five benchmark test sets, yielding 15 points per violin. The horizontal line denotes the median. Three stratification levels are used: High Potency (MIC *<* 10 *µ*M), Intermediate Potency (10 *µ*M ≤ MIC *<* 100 *µ*M), and Low Potency (MIC ≥ 100 *µ*M). A consistent degradation of correlation is observed as peptide potency increases. **b:** Violin plots with overlaid swarm plots representing the same stratified correlation scores, but where both training and evaluation are restricted to the corresponding MIC stratum — that is, each model is retrained on the subset of training data matching the potency threshold. Notably, the High Potency stratum retains the largest training set, followed by Intermediate Potency and Low Potency. Results aggregate all three models across five benchmark splits, yielding 15 points per violin. Despite this data enrichment strategy, correlation performance continues to degrade with increasing peptide potency.

